# Grain Utilization by the Gut Microbiome as a Human Health Phenotype to Identify Multiple Effect Loci in Genome-Wide Association Studies of *Sorghum bicolor*

**DOI:** 10.1101/2023.09.20.558616

**Authors:** Nate Korth, Qinnan Yang, Mallory J. Van Haute, Michael C. Tross, Bo Peng, Nikee Shrestha, Mackenzie Zwiener, Ravi V. Mural, James C. Schnable, Andrew K. Benson

## Abstract

A growing epidemic of complex lifestyle diseases such as obesity and metabolic diseases are explained in part by dysbiosis of the human gut microbiome. The gut microbiome, comprising trillions of microorganisms, contributes to functions ranging from digestion to the immune system. Diet plays a critical role in determining the species composition and functionality of the gut microbiome. Substantial functional metabolic diversity exists within the cultivated grain crops which directly or indirectly provide more than half of all calories consumed by humans around the globe, however much of this diversity is poorly characterized and the effects of such diversity on the human gut microbiome is not well studied. We employed a quantitative genetics approach to identify genetic variants in sorghum that alter the composition and function of human gut microbes. Using an automated high-throughput phenotyping method based on *in vitro* microbiome fermentation of grain from a diverse population of *Sorghum bicolor* cultivars, we demonstrate sorghum genetics can explain effects of grain variation on fermentation patterns of bacterial taxa across multiple human microbiomes. In a genome-wide analysis using a sorghum association panel, we identified fifteen multiple-effect loci (MEL) where different alleles in the sorghum genome produced changes in seed that affect the abundance of multiple bacterial taxa across two human microbiomes in automated in vitro fermentations. In a number of cases parallel genome-wide association studies conducted for biochemical and agronomic traits identified seed traits potentially causal for the link between sorghum genetics and human microbiome outcomes. This work demonstrates that genetic factors affecting sorghum seed can drive significant effects on human gut microbes, particularly bacterial taxa considered beneficial. Understanding these relationships will enable targeted crop breeding strategies to improve human health through gut microbiome modulation.

## Introduction

The gut microbiome is a complex ecosystem of microorganisms residing in the gastrointestinal tract. Species composition and function of the gut microbiome have emerged as crucial determinants of both human health and predisposition to disease, contributing to the metabolism of nutrients, the synthesis of vitamins, the development of the immune system, the maintenance of the intestinal barrier, and the protection against pathogens. Dysbiosis, imbalances, or abnormal configurations in the composition and function of the gut microbiome has been linked to a number of health conditions in humans, including inflammatory bowel disease, diabetes, metabolic disorders, heart disease, and even mental health outcomes^1–4^. While several factors shape species composition and function of the human gut microbiome, diet is one of the most significant determinants of both the abundance of different specific microbial taxa within the gut and overall human gut microbiome diversity^5,6^. Different foods vary in the presence, absence, or content of dietary components that can each promote or inhibit the growth of specific bacterial taxa. The content and composition of digestion-resistant starches, fibers, and polyphenols found in food have been shown to have extensive impacts on gut microbes and human health in *in vitro* and *in vivo* systems^7–9^.

Efforts to understand and harness the role diet plays in shaping the function and composition of the human gut microbiome have historically focused on different diets (e.g. Western vs Mediterranean) or have evaluated the effect of adding a known or suspected prebiotic compound, such as purified starches or fibers to an existing diet^10–12^. Comparatively less effort has been made to investigate the impact of genetic and metabolic diversity within individual food crops on the human gut microbiome. However, our data from our recent in vitro microbiome fermentation studies with grain from functional genetic variants that affect starch composition (waxy) and seed protein composition (opaque-2) in individual varieties of sorghum and maize respectively have shown such variants can produce dramatic changes in the composition and function of human gut microbiomes^13–15^. We have also used quantitative genetics techniques to understand how plant genetic variation influences the human gut microbiome^15^. This is a powerful strategy that presents the opportunity to discover and characterize previously unknown genetic loci controlling dietary components (seed components) that can cause quantitative changes in microbiome fermentation patterns, including previously uncharacterized links between known dietary components and specific bacterial taxa^15^.

Substantial genetic and metabolite diversity exists within the primary gene pools of many of the most widely cultivated and consumed crops around the globe. Amid such diversity, segregating genetic variants within crop species can control not only agronomic traits of the plant, but also control the presence or relative abundance of bioactive compounds that can influence the composition of the human gut microbiome. In the case of sorghum (*Sorghum bicolor*), a grain closely related to maize that acts as a dietary staple and key component of food security in numerous parts of Africa and south Asia, substantial variation exists in the quantity and identity of complex fibers, resistant starch, and phenolic compounds – including condensed tannins^16,17^. Here we employed a sorghum diversity panel representative of global sorghum genetic and phenotypic diversity^18,19^ and a high-throughput *in vitro* microbiome fermentation assay^15^ to identify genetic loci in sorghum associated with change in the composition or function of gut microbiomes from different human donors.

## Materials and Methods

### Plant germplasm and Growth Conditions

The seed for the Sorghum Association Panel (SAP)^18^ used in this study was generated in a field increase conducted in 2020 at the University of Nebraska-Lincoln’s Havelock farm (N° 40.861, W° 96.598). While this field experiment has been previously described in detail^20^, briefly 344 sorghum genotypes listed in table S2, were planted on June 08, 2020. Each plot consisted of a single 2.3-meter row of plants of a single genotype with 0.76 meter spacing between parallel and sequential rows. Two to three panicles per plot were randomly selected and hand-harvested on October 16^th^, 2020. Sorghum seed was dried and cleaned before being processed.

### Biochemical analysis of grain samples

The tannin content of each sorghum genotype was quantified using an established vanillin acidification protocol^21^. Briefly, 1 mL of 4% HCl in methanol was added to 25 mg of ground sorghum grain and incubated at 30 ℃ for 20 minutes. Samples were centrifuged and a 100 µL aliquot was added to 500 µL 0.5% vanillin 4% HCl in methanol and a 4% HCl blank and incubated for 20 minutes. Absorbance at 500 nm was measured using a UV/VIS spectrophotometer. Final tannin contents were calculated by subtracting the absorbance of the blank from the absorbance of the sample and vanillin and comparing it to a catechin standard. Tannin content is reported as mg catechin equivalent lines. Lines that contained less than 1 mg catechin equivalent tannins were classified as “non-tannin” lines in downstream analysis.

Seed protein, oil, and starch concentrations were estimated using near-infrared reflectance (NIR). Each line was dried, cleaned, and then scanned using a Perten DA 7250 NIR spectrometer (Perten Instruments, Hägersten, Sweden). A random sample of the seed harvested from the selected panicles was scanned on the instrument using the small sized sample tray. This measurement procedure was repeated five times per plot with different samples of grain (sampled with replacement). Scans that produced negative estimated values for any grain component were dropped. For each of seed protein, oil, and starch, the median value across all remaining samples within a genotype/line was used for downstream analysis.

Data describing sorghum seed color was collected using a pre-trained MaskR-CNN model on rice seeds to detect and segment the seeds from sorghum seed scans obtained using EPSON scanner^22^. Once the model performed pixel level segmentation for region of interest (ROI); seed regions in the scanned images, three-pixel values; red, green, and blue channels were extracted for each ROIs using custom Python code. For each color channel in a seed scan, the pixel values across all ROIs were averaged. This averaged RGB data as well as 3 principal components calculated using the prcomp function in R was utilized for further analysis.

### Collection of publicly available SAP phenotypes

In addition to the newly generated sorghum phenotypes generated above, a set of 172 trait datasets mined from previously published studies of the sorghum association panel^23^ were included in the genome wide association studies to identify potential overlap between genes/genomic loci influencing directly observable sorghum traits and those influencing variation in the human gut microbiome (Table S3).

### Automated *in vitro* microbiome screening (AiMs) processing, sequencing, and short chain fatty acid (SCFA) analysis

Fecal sample-derived microbiomes were collected from twelve volunteers; eight fecal microbiomes were analysed in pilot screen, two in mapping experiment, and all twelve in downstream validation. Stool samples were diluted in phosphate buffer saline with 10% glycerol and homogenized using an Interscience BagMixer 400 and filtered with Labplus 6x9 filtra-bags. The filtrate was immediately aliquoted frozen at -80℃ for storage. The University of Nebraska IRB (approval number 20160816311EP) approved the sample collection method. Approximately 3 g of sorghum seed for each of 344 lines was milled in a GenoGrinder 2025 (SPEX SamplePrep, Metuchen, NJ, USA) with 2, 7/16” stainless steel ball bearings in 15 mL polycarbonate vials at 1600 rpm for 6 minutes. Twenty mg (+/-0.05 mg) of milled flour was dispensed into 96-well plates in a randomized complete block design with a Flex Powderdose GDU-p (Chemspeed Technologies AG, Füllinsdorf, Switzerland). Each of the three blocks consisted of four, 96-well plates, with each line of sorghum present in at least one well in each block. Sorghum flour was digested following established protocols^15^. Sorghum flour was hydrated in 425 µl water and cooked in boiling water for 20 minutes, agitating at 30-second intervals. Samples were treated with 45 µl of 500 mM HCl and 10% pepsin (P7000; Sigma, 470 St. Louis, MO) at 37 °C for 1 hour. The small intestinal phase was initiated by adding sodium maleate buffer (pH = 6, containing 1 mM CaCl2) and NaHCO3. Pancreatin (P7545; Sigma, St. Louis, MO) with amyl glucosidase (E-AMGDF, 473 3,260 U/mL, Megazyme) was added before incubation at 37 °C for 6 hours. Post digestion, samples were transferred to 96-well dialysis plates (MWCO 1,000; 475 DispoDialyzer; Harvard Apparatus, Holliston, MA, USA) and dialyzed in five gallons of distilled water replaced at 12-hour intervals for 72 hours at 4 °C with agitation to facilitate the removal of small molecules. The digested and dialyzed sorghum flour was transferred to 1 mL 96-well plates. Fifty microliters of fecal microbiomes and 50 microliters of 10x fermentation media containing 1 g Bacto casitone, 1 g yeast extract, 2 g K2HPO4, 3.2 g NaHCO3, 3.5 g NaCl, 1 mL hemin solution (KOH 0.28 g, 95% Ethanol 25 mL, hemin 100 mg and ddH2O to 100 mL), 0.05 g bile salts, 0.5 g/L cysteine HCl, 0.6 mL resazurin (0.1%), 10 mL ATCC trace mineral supplement, 3.6 mL VFA solution (17 mL acetic acid, 1 mL n-valeric acid, 1 mL iso-valeric acid, 1 mL iso-butyric acid mixed with 20 mL of 10 mM NaOH), 10 mL ATCC vitamin supplement and 1 mL vitamin K-3 solution (0.14 g vitamin K-3 in 100 mL 95% ethanol)^24^ reduced by the addition of 50% Oxyrase (SAE0013; Sigma, St. Louis, MO) were added in an anaerobic chamber. Fermentations were incubated for 24 hours at 37 ℃. Samples were centrifuged, bacterial pellets were used to determine microbiome composition and the supernatant was retained for SCFA analysis. Both pellets and supernatant were stored at -80 ℃ until processing.

DNA was extracted from bacterial pellets in the BioSprint 96 workstation (Qiagen, Germantown, MD) and the BioSprint 96 one-for-all Vet Kit utilizing ASL buffer (19082; Qiagen, Germantown, MD) and bead beating. Sequencing of the V4 region of the 16S rRNA gene was achieved by amplification and indexing by PCR primers as previously described^25^. Libraries were normalized using a SequalPrep Normalization Plate (96) Kit (A1051001; Invitrogen, Frederick, MD). Paired-end sequencing was performed on an Illumina MiSeq^26^ at the Nebraska Food for Health Center. An average of 31,664 reads per sample was obtained.

Dada2 within the QIIME2 program^27^ was employed to extract amplicon sequence variants (ASVs) and assign each taxonomy based on the 132^nd^ release of the SILVA 16S reference database^28^. Forward and reverse reads were trimmed to 220 and 160 bp respectively to remove low-quality sequence data. ASVs present in only a single sample or composed of less than 15 reads total were removed from the analysis as were all microbial taxa at family, genus, and ASV levels with less than five reads in seventy-five percent of the samples. Baseline microbiome samples and microbes fermented in media only (absent of sorghum) were collected as controls.

### Quantification of Faecalibacterium prausnitzii

The abundance of *F. prausnitzii* was measured using a quantitative PCR utilizing primers: FprauF: TGAGGAACCTGCCTCAAAGA; FprauR: GACGCGAGGCCATCTCA developed by Linstad et al.^29^. The qPCR reactions, conditions, and standard curve calculation were as previously reported^15^.

### Variance Partitioning Analysis

The responses of gut microbiomes of eight human donors to a subset of 24 sorghum genotypes listed in table S2 were evaluated. For each ASV observed in at least 20% of samples, variance was partitioned using a linear mixed model using the relative abundance of a single genus of bacteria for a given subject as a response and fitting sorghum genotype, color, and tannin content as random effects in the Sommer package in R^30^ . Information on sorghum color and condensed tannin content was collected from publicly available data^31^.

### Detection and Removal of Outliers Among Technical Replicates

A three-stage outlier detection and removal procedure was employed for microbiome traits prior to quantitative genetic analysis. In the first stage, the Jaccard index of beta diversity was calculated for each sample. For technical replicates, defined as samples generated using the same sorghum grain sample and treated with the same human gut microbiome, any single sample which was greater than 0.2 from the midpoint of all technical replicates for that combination of sorghum genotype and human gut microbiome were removed. In the second stage, the mean and standard deviation for each microbial trait, including diversity, relative abundance, SCFA, and polymicrobial traits were calculated. Individual values greater than 5 standard deviations from the mean were dropped, while other values from the same technical replicate were retained in the analysis. In the third stage: within replicates, an upper and lower threshold was set by calculating the upper and lower quantiles of the data and adding to the upper and subtracting from the lower, the interquartile range x 2.5, values outside this range were removed. Across all stages less than one percent of the data was identified as outliers.

### Estimating the Heritability of Microbial Traits

Heritability (proportion of phenotypic variation due to genotypic variation) was estimated using a linear mixed model as implemented in the Sommer package in R^30^. For each microbiome phenotype a linear mixed model was fit including sequencing batch, plate number, and sorghum genotype identity as random effects. Reported heritability values are the quotients of variance in the response explained by sorghum genotype divided by the sum of the variance in the response explained by sorghum genotype and residual error.

### Best linear unbiased estimates calculation and estimates of polymicrobial traits

A linear mixed model was fit to each trait using the Sommer package^30^ in R to calculate best linear unbiased estimates (BLUEs) . Sorghum genotype was fit as a fixed effect. Sequencing batch, plate, row, and column were fit as random effects. The Sommer function predict.mmer was used to calculate BLUEs based on the mixed model.

### Polymicrobial traits calculated from 16s rRNA sequencing data

One hundred principal components (PCs) summarizing a matrix of the BLUE values for the 50 ASVs with the highest estimated heritabilities from each subject (100 ASVs in total) were calculated using the prcomp function in R with the center and scale parameters set to true generating 100 PC scores for each sorghum genotype.

Four estimated prebiotic potential index (PPI) values were calculated using the following equations. The first value was calculated based on the equation: PPIa, the sum of beneficial microbes minus detrimental bacteria, and is denoted as PreInd. The second value based on PPIb is transformed based on the difference of each bacterium incorporated to their mean value and is denoted as PreIndT. Both equations were also applied using only beneficial microbes and are denoted as PreIndB (non-transformed) and PreIndBT (transformed). A list of organisms accounted for by the prebiotic index is reported in supplemental table 1.

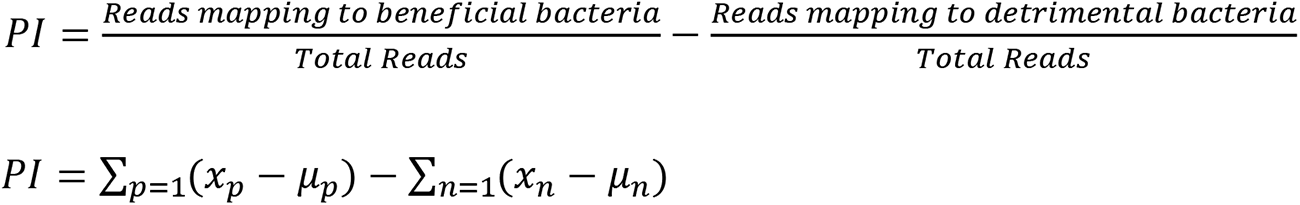

Where x*_p_* and x*_n_* represent the relative abundance of a given beneficial or detrimental microbe respectively and *µ* represents the mean value of that microbe. Prebiotic index correlated strongly with butyrate production in both human subjects (SFigure 1).

An autoencoder neural network was employed to extract 10 latent variables describing sample to sample variation in patterns of microbial abundance. As separate sets of taxa were present in each subject, separate encoder and decoder architectures were trained on datasets from subject 1 and subject 2. Encoders and decoders were trained on the raw data (e.g., before consolidating technical replicates) after the detection and removal of outlier samples. The dataset for subject 1 comprised 1073 samples described by 99 heritable microbiome traits. The dataset for subject 2 comprised 1075 samples described by 106 heritable microbiome traits. Missing values for individual traits were imputed using the median value for the same trait among other samples which contained the same grain from the same sorghum genotype as the missing sample. In each case, the data was split at a ratio of 4:1 into training and validation sets.

The auto-encoder model was implemented using PyTorch (v1.10.2)^32^ package and the python (v3.9) programming language^33^. The model consisted of an encoder containing an input layer, three hidden layers, and an output layer as well as a decoder also containing an input layer, three hidden layers, and an output layer. For subject 1, the layers consisted of 99, 220, 300, 224, and 10 neurons in the encoder and 10, 224, 300, 220, and 99 neurons in the decoder. For subject 2, the layers consisted of 106, 220, 300, 224, and 10 neurons in the encoder and 10, 224, 300, 220, and 106 neurons in the decoder. The number of neurons in the first layer of the encoder and final layer of the decoder was set equal to the number of microbiome traits describing each subject. In the final layer of the decoder a hyperbolic tangent activation function was employed while for all other layers the activation function used was the scaled exponential linear activation unit. The model was trained for 1,000 epochs using a mean absolute error loss function and a standard gradient descent optimization algorithm at a learning rate of 0.1. The best model was saved based on the lowest reconstruction loss on the validation dataset after each epoch. Final trained encoders were used to summarize the variance in each technical replicate. BLUEs for each of the 10 autoencoder derived latent variables derived from each subject (20 total latent variables) were calculated as described above.

### Genome-wide association study

A marker set consisting of approximately 43 million genetic markers (e.g. SNPs, indels, and structural variants) derived from whole genome resequencing of the sorghum panel^19^ were used in GWAS analysis. Sorghum accessions phenotyped in this study and not resequenced as part of Boatwright et al., were removed, reducing the number of genotypes to 340. Markers with a minor allele frequency of less than 0.1 and markers scored as heterozygous in more than 10% of sorghum lines genotyped were removed, resulting in a final set of 4,090,874 markers employed for GWAS. For each trait, including agronomic traits from Mural et al., and BLUEs from traits measured in this study (e.g. tannin concentration, seed composition, seed color, and microbial traits) GWAS was conducted using the FarmCPU algorithm^34^ as implemented in rMVP^35^. For each trait the first three principal components of genetic marker variation – calculated using rMVP -- were included as covariates. Kinship was controlled for as described in the FarmCPU algorithm. Resampling model inclusion probability (RMIP) was calculated by analysing each trait 100 times, each with a random 10% of the data masked^36^. Markers were considered significant if they had a p-value less than 0.05/ 861,521.36, a threshold set by a Bonferroni correction using the effective number of markers calculated by the genetic type 1 error calculator, GEC^37^. Genetic markers were considered significant if they were detected with a p-value more significant than the significance threshold in at least 10 of the 100 FarmCPU iterations corresponding to an RMIP threshold of 0.1.

### Defining Major Effect Loci (MEL)

To identify regions potentially harbouring allelic variants with major effects on the human gut microbiome, each chromosome was split into 75 equally sized bins. As the sizes of each sorghum chromosome are distinct, this division method produced bin sizes ranging from 0.79 to 1.07 MB for different chromosomes. A thresholding strategy was used to identify bins containing significant GWAS hits (defined as an RMIP value >= 10) for at least five microbial traits derived from each of the two human subjects. Hits for additional traits present in genomic bins adjacent to those selected for manual examination were merged as appropriate to control for the arbitrary placement of bin boundaries. Borders of bins meeting the threshold were then adjusted around the most frequent significant marker in the bin (marker affecting the most traits) using linkage disequilibrium (LD) of genetic markers within 100 KB of the most frequent significant using the TASSEL program^38^ to assist in defining the outer edges of the candidate gene space surrounding a given cluster of linked GWAS hits. Markers in high LD (r^2^>0.5) with significant Markers used to define linkage blocks constituting final MEL regions. Linkage disequilibrium among all Markers identified as significantly linked to at least one microbiome trait within a given MEL was used to categorize MELs into those likely to result from a single causal variant (high, maximum r^2^>0.5 and the average r^2^>0.05) those likely to result from the colocalization of two more independent variants (split maximum r^2^>0.5 and the average r^2^<0.05, low maximum ^r2^ <0.5 linkage).

### Candidate Gene Detection

A list of genes within the boundary of a MEL was compiled based on the gff3 gene annotation file for the v3.1 build of the sorghum genome^39^. If less than five genes were present within the boundaries of a MEL, the window was expanded by 50 kb in both directions. Transcriptional data from sorghum five and ten days after pollination, and embryo and endosperm twenty-five days after pollination was obtained from publicly available data^40^ and used to highlight genes that are expressed in the seed and filter out genes not expressed in seed. Genes where all seed expression values (FPKM) were less than 1, were removed from further analysis.

### Validation of MEL6A haplotype

Sorghum genotypes were pooled into four categories based on tannin content and allele at the SNP associated with the most microbial metrics in MEL6A (S06_43330339). Two pools of each of the four groups were made from ten randomly selected sorghum genotypes, 2.5 g of each pool was processed in the same digestion protocol described above. Fermentations were conducted *in vitro* using the fecal microbiomes of twelve human subjects collected as previously described on the sorghum pools. DNA extraction, 16S amplicon sequencing, and qPCR were conducted as described above.

### Statistical Methods

Data processing, compilation of diversity metrics, PERMANOVA, Wilcoxon tests, PCA analysis, BLUE calculation, FarmCPU GWAS, SNP binning, and data visualization were conducted in the R statistical framework v4.0.2^41^ utilizing the packages Sommer v4.1^30^, phyloseq v1.42.0^42^, Metagenomeseq v1.4^43^, tidyverse v2.0^44^, rMVP v1.0.6^35^, circlize^45^, and ggplot2 v3.4.2^46^. BLASTp, hosted on National Center for Biotechnology Information web service^47,48^ was used to align protein sequences and calculate sequence similarly. LefSE analysis was performed in the galaxy web application^49,50^. Gene functional annotations and analysis of gene sequence variation were based on the v3.1.1 sorghum reference database^40^ accessed via the Phytozome database^39^. The InterPro database within Phytozome was used to assign functional annotations to candidate genes. R code to conduct the analyses described in this paper is available at https://github.com/natekorth/SAP.

## Results

Variation among sorghum genotypes explains substantial variation in human gut microbiomes Seed from 344 lines of the sorghum association panel grown in Nebraska in 2020 was screened across the microbiomes of two human subjects in a replicated *in vitro* fermentation experiment. The gut microbiomes of two human subjects were selected from a set of eight candidate subjects. These two microbiomes were selected based on baseline composition and how microbial communities responded to *in vitro* fermentation with grain samples from a subset of 24 sorghum lines selected from the SAP. A variance partitioning analysis determined that the microbiomes of the two selected subjects (S766 and S777) included the largest number of microbial taxa where more than 20% of total variance could be attributed to differences between sorghum genotype (Table S4). The two selected microbiomes are referred to as subject 1 (S766) and subject 2 (S777) below. Notably, tannin content in the sorghum lines explained much of the variation in the abundance of different microbial taxa present in subject 1 whereas the microbiome of subject 2 was substantially less tannin responsive, implying that the nutritional basis for different fermentation patterns across genotypes was due to different components of the seed and/or differences in species and strain presence/absence in one or both microbiomes.

AiMS-based phenotyping of the entire SAP resulted in a total of 2,211 individual in vitro fermentations (1,106 from the subject 1 microbiome and 1,105 from the subject 2 microbiome, excluding one sample lost during processing). Changes in the abundance of microbial taxa post-fermentation were assayed via 16S sequencing, and metabolic outcomes of fermentation were assayed using gas chromatography. A three-step outlier identification process resulted in the removal of 28 individual AiMs reactions and 563 individual trait value observations for subject 1 and 26 samples and 650 individual trait values for subject 2. The microbiome of subject 1 comprised of 3,130 individual ASVs and subject 2 comprised of 2,368 ASVs. ASVs not present (at least 5 reads) in 75% of the samples from a single subject were not considered traits in downstream analyses. An estimate of heritability for each taxon was calculated and used to remove traits where variation could not be explained by variation in the genotype (h^2^<0.05).

After filtering by abundance and heritability, the microbiome of subject 1 was comprised of 71 ASVs collapsed into 48 genera and 21 families and the microbiome of subject 2 was comprised of 76 ASVs, collapsed into 50 genera and 19 families. Five different metrics of alpha diversity (Observed ASVs, Shannon, Simpson, Inverse Simpson, and Fisher) and twenty-two additional metrics that reduce the dimensionality to describe each microbiome (referred to as polymicrobial traits) were also included (ten PCs, ten latent variables from the autoencoder analysis, and two versions of the prebiotic index outlined in materials and methods). We also used metabolic end-products of microbial fermentation (short chain fatty acids and branched-chain fatty acids) measured post fermentation from both subjects as quantitative traits. A prebiotic index calculated from 16S abundance data of each AIMS reaction, exhibited roughly equivalent distributions across between the two subjects (Figure 1A). In contrast, butyrate production, a microbial fermentation end produce assayed from the same AiMS reactions showed quite different distributions between the microbiomes, illustrating distinctive effects of sorghum genotype on fermentation patterns of the two microbiomes (Figure 2A).

**Figure 1.**
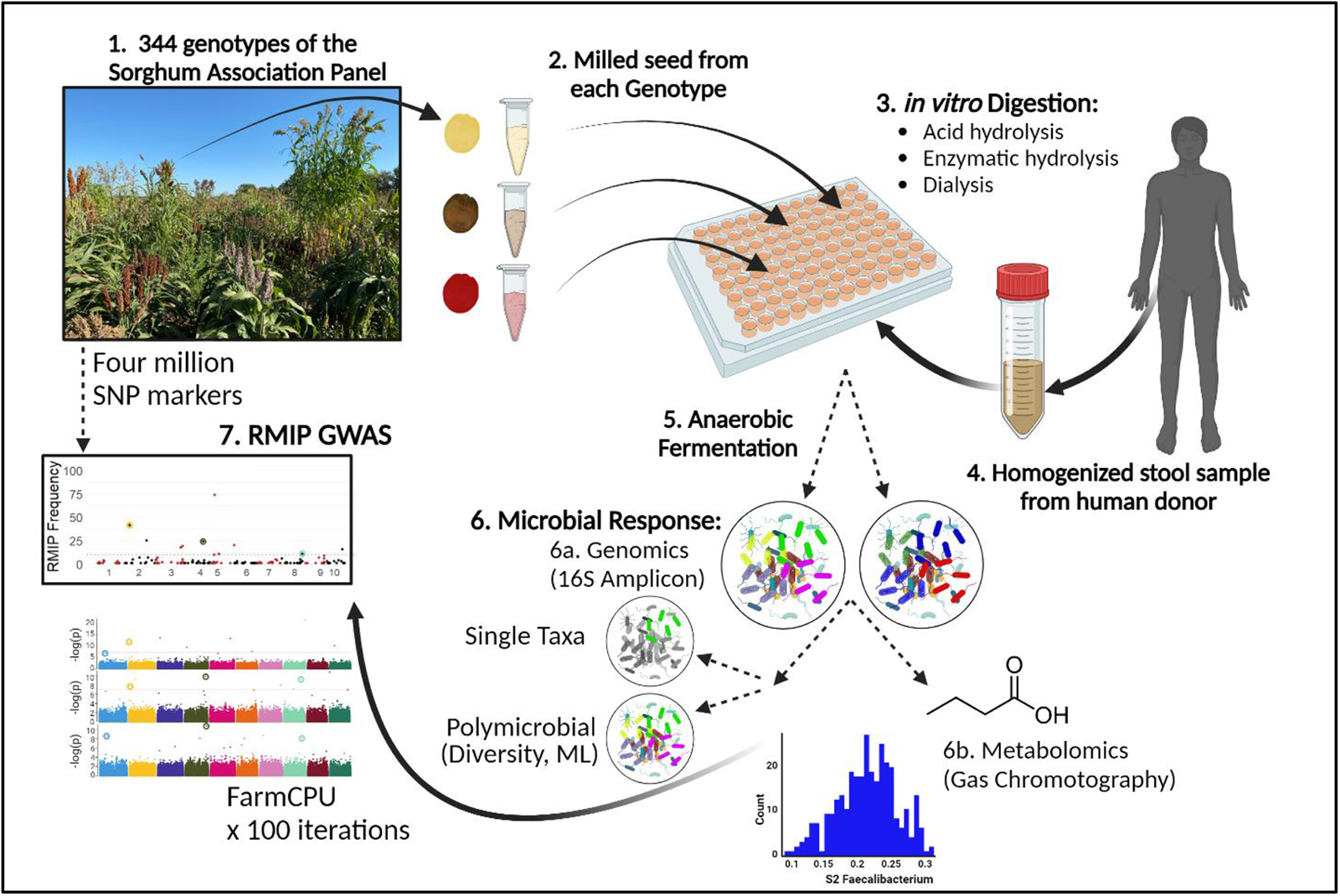
Graphical abstract describing the Automated *in vitro* Microbiome screening (AiMs) platform. Varieties of the SAP, grown in Nebraska in 2020 were collected and ground into flour (1 and 2). Following the dispensing of each flour into 96-well plates, a simulated human digestion protocol including mechanical, chemical, and enzymatic treatments was applied to each sample. Small molecules were removed in a dialysis protocol simulating absorption in the small intestine (3). The digested material was fermented with human gut microbes collected from human stools in anaerobic conditions (4 and 5). Sequencing and metabolomics were utilized to measure the microbial response to each sorghum genotype (6). All metrics describing the microbiome response was treated as a phenotype of sorghum in a genetic mapping study using a resampling method of FarmCPU (7). This Diagram was created with BioRender.com.

A total of 127 traits describing the microbiome of subject 1 and 129 traits describing the microbiome of subject 2 where at least 5% of total variance was explained by differences between sorghum genotypes in a variance partitioning analysis (Figure 2B and Table S4) were selected for quantitative genetic analysis. The abundance of many microbes in fermented samples were correlated with variation in the sorghum seed traits, including traits that would be expected to impact specific microbial taxa and groups of microbial taxa. For example, observed expected relationships between fiber-fermenting organisms comprising the Prebiotic Index of Subject 2 and seed fiber content (Figure 2C – top). We also detected significant relationships between *Faecalibacterium* from Subject 1 and condensed tannins (Figure 2C – middle) that were observed in a recent genetic analysis by^15^. In addition, there is correlation between *Akkermansia* from Subject 2 and oil content (Figure 2C-bottom) that aligns with known utilization of unsaturated fatty acids by *Akkermansia*^51^.

In many cases the same sorghum lines produced similar functional impacts – as summarized via a prebiotic index -- on polymicrobial metrics of the subject 1 and subject 2 microbiomes (Figure 2D), despite the large baseline differences present between the two microbiomes (Figure S2). Best linear unbiased estimators (BLUEs) were calculated for each microbiome metric and are available in supplementary dataset S1.

**Figure2.**
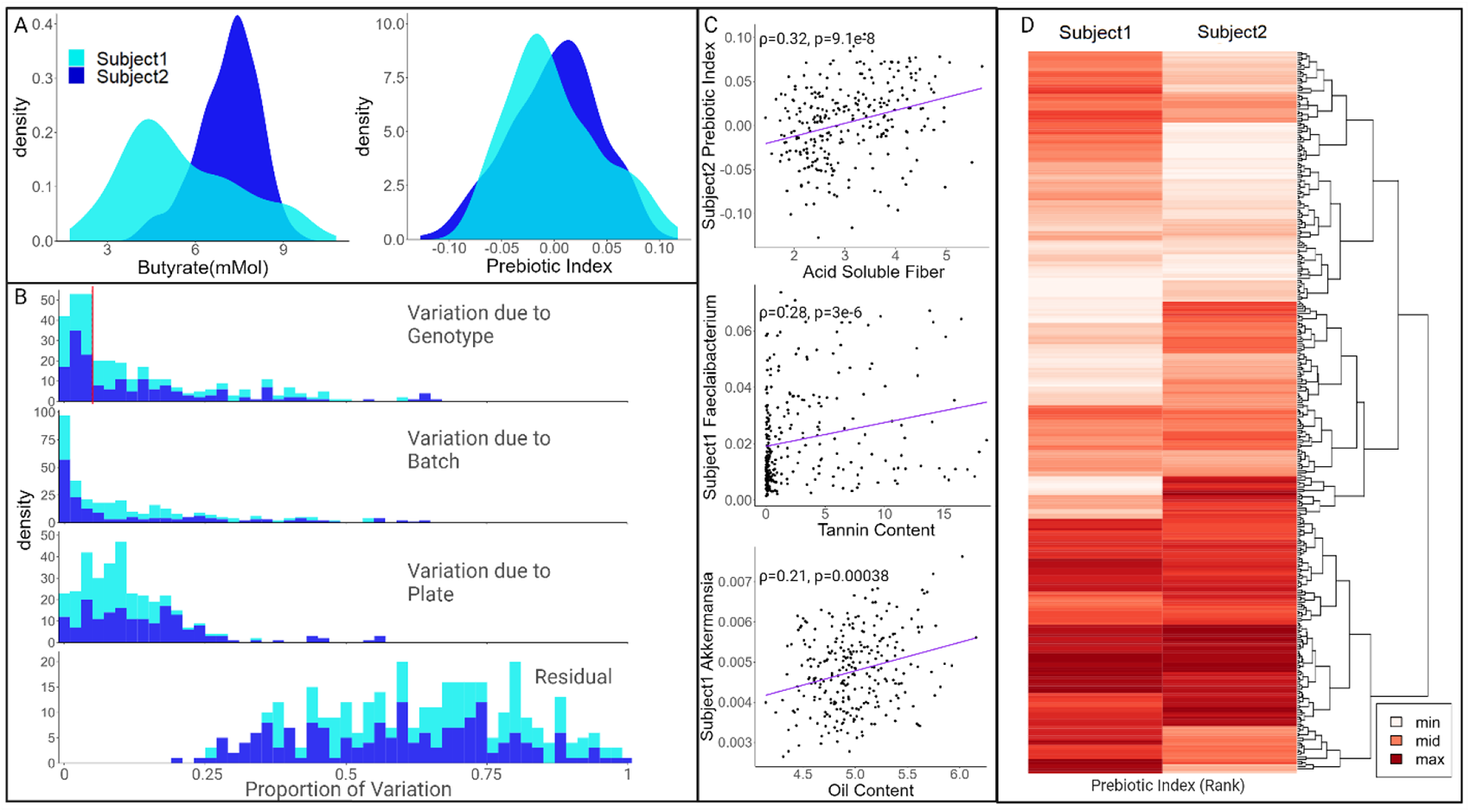
Sorghum genetics impacts microbial utilization of sorghum seed. A.) Distributions of BLUEs calculated for butyrate concentrations and prebiotic indices for 340 sorghum genotypes within microbiomes from two human subjects. B.) Variance partitioning of 342 microbiome metrics indicates that for many metrics, variation in the microbiome can be explained by variation in sorghum genotype. C.) Spearman correlation analyses of microbiome metrics and sorghum seed composition traits are in line with hypotheses based on existing literature. D.) Ranked sum abundance of prebiotic index in the microbiomes of two human subjects reveals similarities between microbial responses in distinct microbial communities.

### GWAS identifies loci in sorghum genome associated with multiple changes in the composition and function of human gut microbiomes

Substantial number of trait associated genetic markers were linked to variation in the abundance of individual microbial taxa, polymicrobial traits, microbial population diversity metrics, and metabolic features (SCFAs) in each subject (RMIP ≥ 10, FarmCPU resampling method). In subject 1, a total of 224 markers were associated with one or more trait and in subject 2, a total of 329 markers were significantly associated with one or more trait (Table 1).

**Table 1.** Number of significant associations in the sorghum genome identified for different categories human microbiome characteristics.

| Microbiome Metric | Subject 1 | Subject 2 |
| --- | --- | --- |
| Single Taxa | 145 | 223 |
| Diversity Metrics | 20 | 15 |
| PolyMicrobial Traits | 25 | 83 |
| Short Chain Fatty Acids | 34 | 8 |

Given the large number of associations, we focused on regions of the sorghum genome containing genetic markers significantly linked to at least ten microbiome metrics with at least one metric from both subjects. These are defined as Multi Effect Loci (MELs) and MELs were identified on nine of the ten *Sorghum bicolor* chromosomes and are numbered based on which chromosome they are found (Table 2). In cases where there are multiple MEL per chromosome, they are assigned a letter A-C based on marker position in ascending order. Linkage analysis among trait associated genetic markers was used to distinguish MELs likely to correspond to a single major causal variant impacting multiple phenotypes vs two or more independent variants located within the same genomic interval. Three, seven, and five MELs were classified as exhibiting low, split, or high linkage respectively (Figure 3). Values for LD in each MEL region can be found in Table S6 and Figure S3.

**Figure 3.**
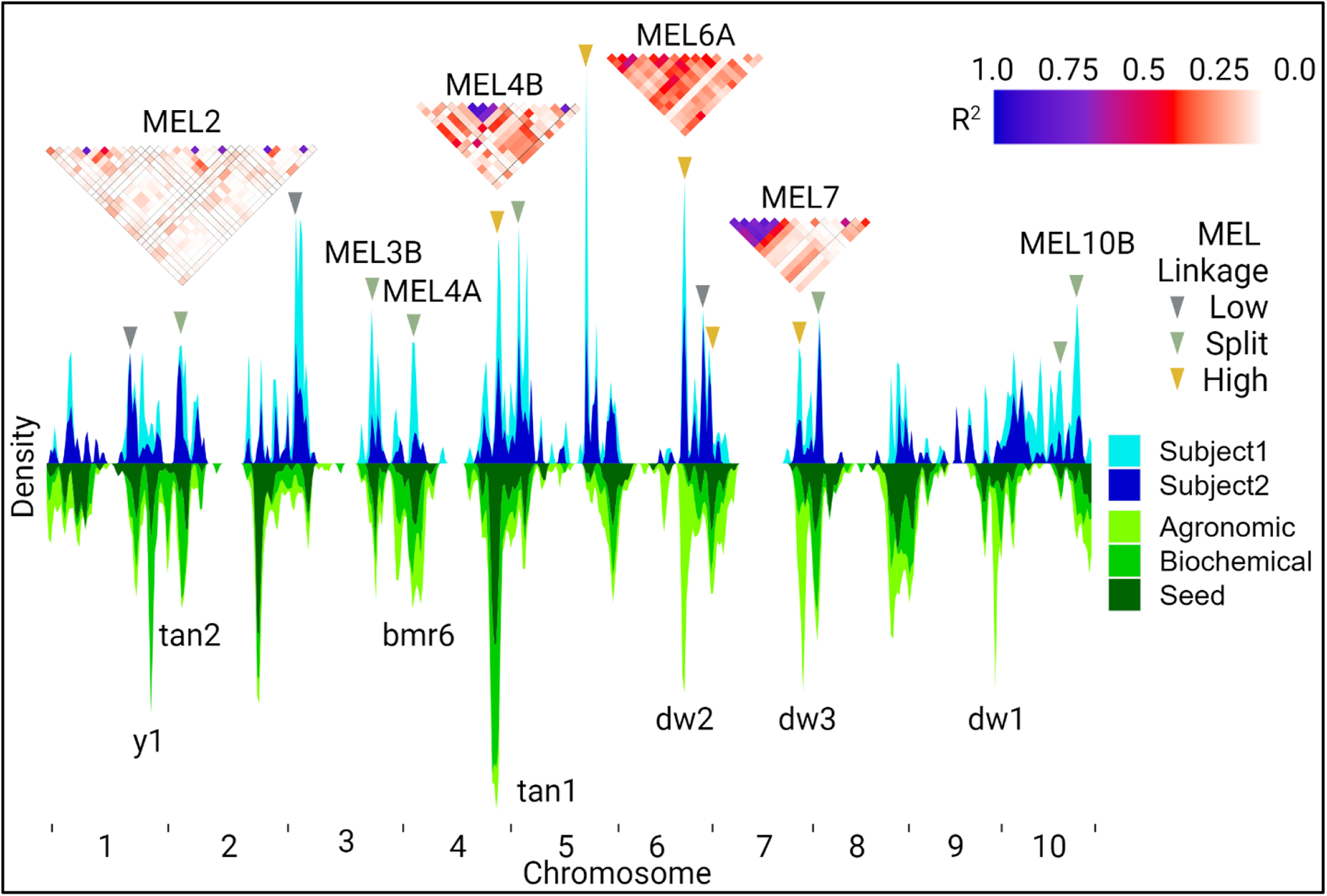
GWAS results in the form of a kernel density plot where the y-axis indicates the frequency of traits associated with a physical position in the sorghum genome. Microbiome metrics have been grouped and colored based on subject (above the center line y-axis) and type of agronomic trait (below the center line y-axis). Linkage between significant markers is shown for four MEL and degree of linkage is denoted by colored triangle heatmaps. The x-axis is ordered by marker position and numbered by corresponding sorghum chromosome.

A large number of phenotypes have been scored from the Sorghum Association Panel^23^. An analysis using a combination of published trait data and new human gut microbiome phenotypes collected as part of this study identified a total of 555, 350, and 426 genetic markers in the sorghum genome associated with one or more agronomic, biochemical, and seed composition traits respectively when GWAS conducted using the same methodology and marker set employed for human microbiome traits. In a number of cases the genetic markers associated with non-microbiome sorghum traits clustered around the locations of previously cloned and characterized large effect sorghum genes (Figure 3; Table 2; Table S7). In all but one MEL (MEL3A), trait associated markers for phenotypes measured from sorghum plants colocalized within MELs identified for human microbiome traits within gene dense regions far from centromeric regions (Figure 4 and Table S7). For example, MEL2 and MEL4B are adjacent to the tan1 and tan2 genes which are known to regulate the accumulation of condensed tannins in sorghum^52,53^. In addition to direct association with microbial taxa phenotypes, genetic markers within these two MELs were also significantly associated with grain color, phenol abundance, and tannin concentration phenotypes (Table S7). *Faecalibacterium* have been previously shown to accumulate in higher abundance in human microbiome samples fermented with high tannin sorghum^15^ and were among the bacterial taxa from both microbiomes that were significantly associated with the genetic markers of these two MELs.

Additionally, MEL10B encompasses the gene, DGAT1, a diacylglyceroal O-acyltransferase critical in lipid biosynthesis^54,55^. Measurements of seed oil mapped to this locus, as did the genus *Akkermansia* from subject 1. *Akkermansia muciniphila* is a known beneficial microbe that has been shown to be stimulated by unsaturated fatty acids^51^, implying that variation in seed oil composition may be driving variation in the *Akkermansia* abundances in the fermentations.

**Figure 4.**
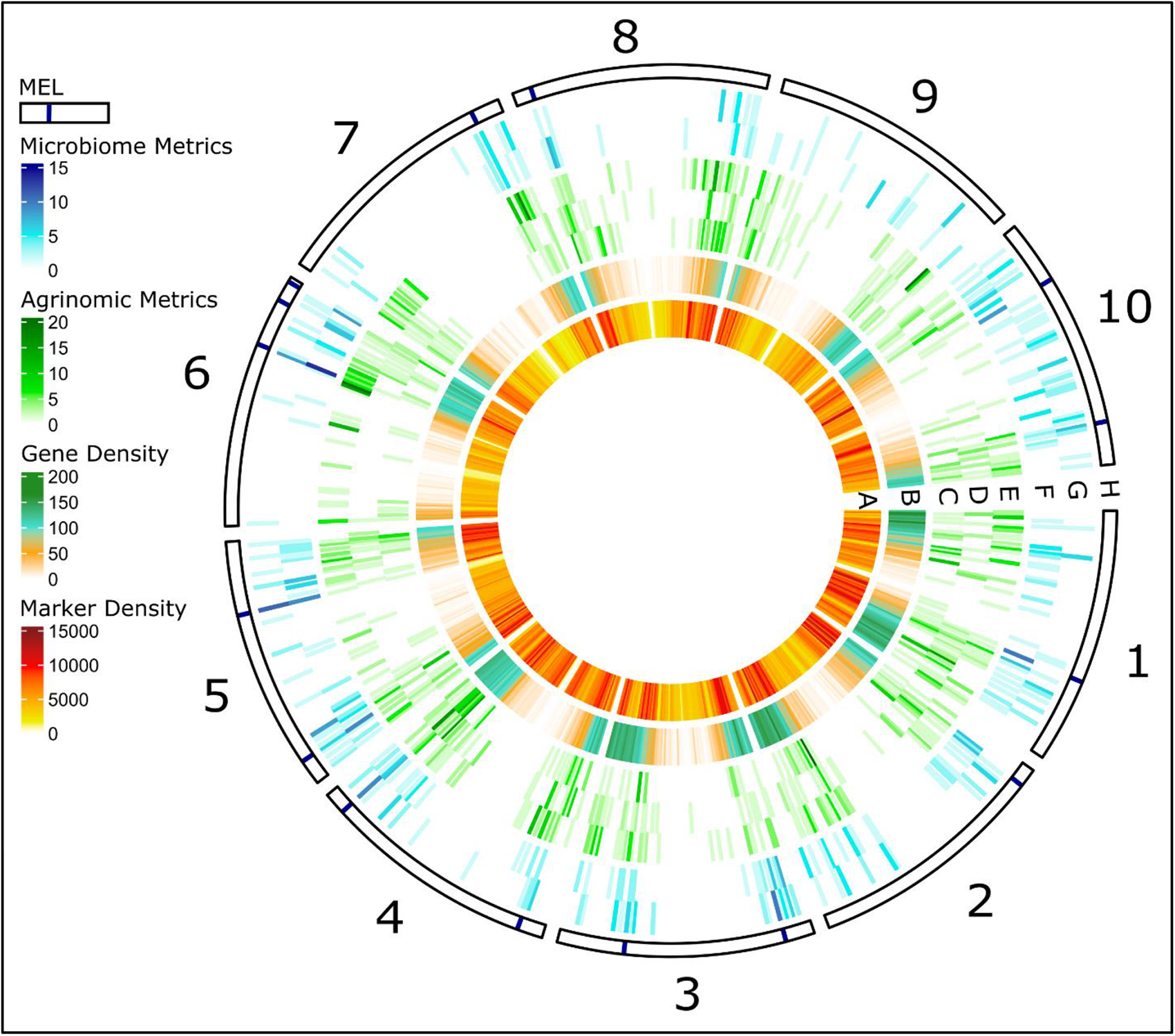
The circular plot illustrates each chromosome split into 75 bin of ∼1MB. Individual tracks depict the SNP density (A), gene density (B), the number of significant markers associated with microbiome metrics from S1 (C) and S2 (D), the number of significant markers associated agronomic (E), biochemical (F), and seed (G) traits, and the locations of MELs (H).

We also identified MELs that were not near known major effect but the co-localized microbiome and agronomic, biochemical, or seed traits (Figure 3) provided some insights as to candidate mechanisms through which sorghum genetics could be affecting the human gut microbiome. For example, in MEL6C, associations with biochemical traits relating to seed phenolics and fiber content were colocalized near significant associations with microbes in the family *Ruminococcaceae* family from both subjects as well as metrics describing the microbial community structure (alpha-diversity and polymicrobial traits). Co-localization of these microbial traits and seed traits suggest that variation at MEL6C could affect the gut microbiome through variation in content of polyphenols, (anthocyanins, condensed tannins), fiber content (e.g. hemicellulose content of cell walls) or an interaction between these two classes of seed components.

**Table 2.**
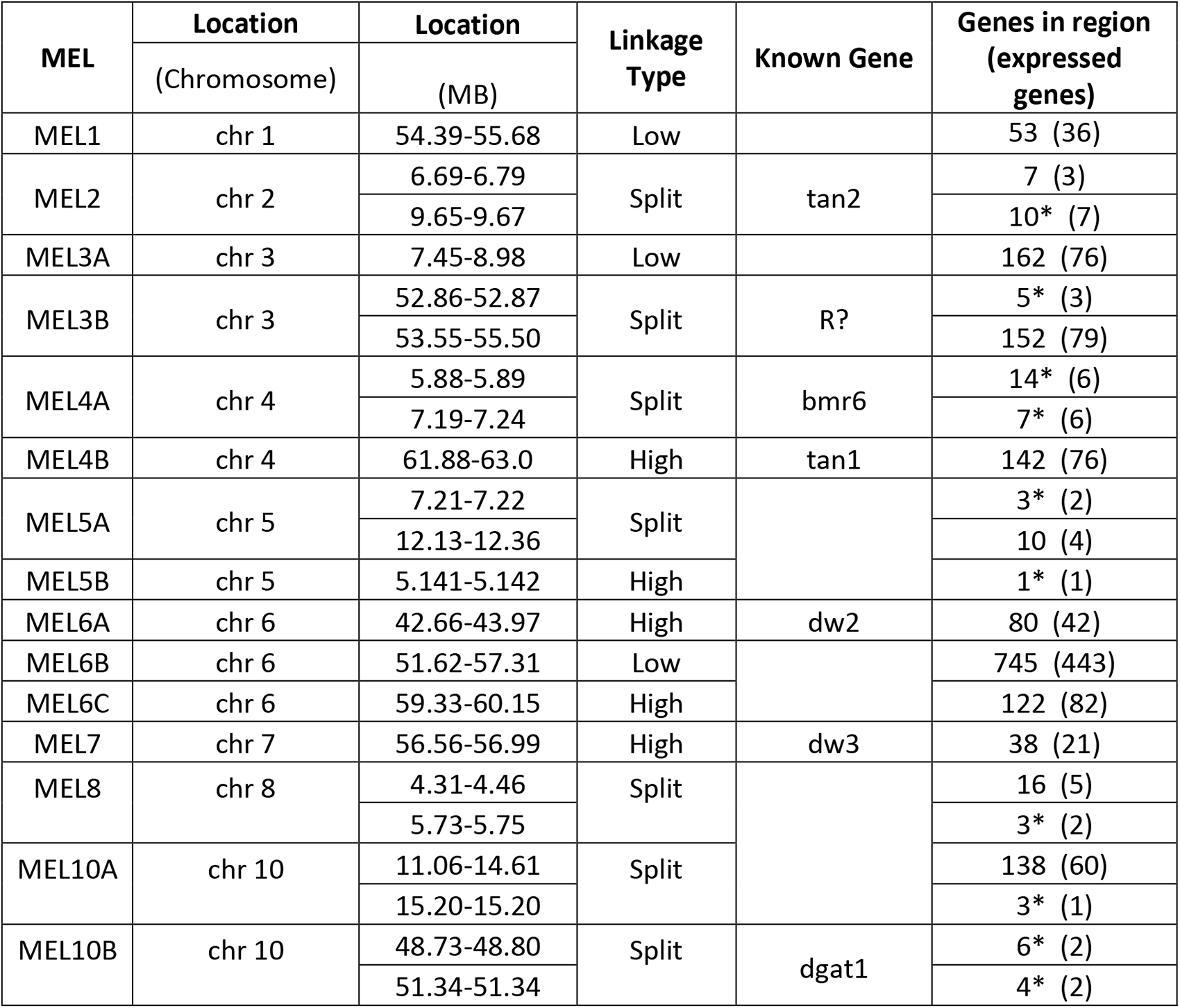
Description of MEL including physical positions two ranges are given to MEL assigned split linkage. Known genes that may be related to seed composition in that region as well as the total number of genes in each region, and how many are expressed in seed. An * denotes cases where no genes were found within the initial MEL region defined by LD and an expanded window (50 additional kilobases on either side of the region) was used to identify potential candidates.

| MEL | Location<br>(Chromosome) | Location<br>(MB) | Linkage<br>Type | Known Gene | Genes in region<br>(expressed<br>genes) |
| --- | --- | --- | --- | --- | --- |
| MEL1 | chr 1 | 54.39-55.68 | Low |  | 53 (36) |
| MEL2 | chr 2 | 6.69-6.79 | Split | tan2 | 7 (3) |
|  |  | 9.65-9.67 |  |  | 10* (7) |
| MEL3A | chr 3 | 7.45-8.98 | Low |  | 162 (76) |
| MEL3B | chr 3 | 52.86-52.87 | Split | R? | 5* (3) |
|  |  | 53.55-55.50 |  |  | 152 (79) |
| MEL4A | chr 4 | 5.88-5.89 | Split | bmr6 | 14* (6) |
|  |  | 7.19-7.24 |  |  | 7* (6) |
| MEL4B | chr 4 | 61.88-63.0 | High | tan1 | 142 (76) |
| MEL5A | chr 5 | 7.21-7.22 | Split |  | 3* (2) |
|  |  | 12.13-12.36 |  |  | 10 (4) |
| MEL5B | chr 5 | 5.141-5.142 | High |  | 1* (1) |
| MEL6A | chr 6 | 42.66-43.97 | High | dw2 | 80 (42) |
| MEL6B | chr 6 | 51.62-57.31 | Low |  | 745 (443) |
| MEL6C | chr 6 | 59.33-60.15 | High |  | 122 (82) |
| MEL7 | chr 7 | 56.56-56.99 | High | dw3 | 38 (21) |
| MEL8 | chr 8 | 4.31-4.46 | Split |  | 16 (5) |
|  |  | 5.73-5.75 |  |  | 3* (2) |
| MEL10A | chr 10 | 11.06-14.61 | Split |  | 138 (60) |
|  |  | 15.20-15.20 |  |  | 3* (1) |
| MEL10B | chr 10 | 48.73-48.80 | Split | dgat1 | 6* (2) |
|  |  | 51.34-51.34 |  |  | 4* (2) |

### Investigation of MEL6A

Among the MELs where we observed co-localization of microbiome and agronomic traits, MEL6A was particularly intriguing as several agronomic and seed traits (seed weight, harvest index, and panicle area) map to the same locus along with a number of individual microbiome traits (*Faecalibacterium* from both subjects, *Escherichia* from subject 1, and *Coprococcus* plus other members of the *Lachnospiraceae* from subject 2) and polymicrobial traits (PC2 in both subjects, LV2 in subject 1 and prebiotic index in subject 2). Centered within MEL6A on the sorghum genome is a tandem array of four cell wall invertase (CWIN) genes located approximately 43 megabases from the start of sorghum chromosome 6 (Figure 5A). Two of the significantly trait associated genetic markers were S06_43329706 and S06_43330339. These markers were immediately adjacent to the tandem array of CWIN genes and they are associated with the most microbial metrics within this MEL, including subject 2 prebiotic index and multiple groups within the families, *Lachnospiraceae*, *Ruminococcaceae*, and *Enterobacteriacae* from both human subjects (Figure 5A). Sorghum has a total of 16 genes annotated as encoding a β-fructofuranosidase, otherwise known as invertase. However only eight of these genes, including four CWIN within MEL6A, are expressed in developing sorghum grain. Moreover, the MEL6A CWIN appear to be exclusively expressed in developing seed whereas the four other seed-expressed invertases are also expressed in leaf and/or root tissue (Figure 5D). The MEL6A CWIN are also part of a six gene sub-family in sorghum that appear to be orthologous to a four gene sub-family of CWIN in Arabidopsis which also exhibits substantial seed-specific expression, the other ten sorghum invertases share more peptide similarity with vacuolar invertases and are likely not cell wall bound. Cell wall invertases in other plant species are known to play important roles in sucrose metabolism, and loss of function mutations of these genes can be pleotropic producing phenotypes influencing the formation of fructooligosaccharides, flower and seed development, seed size, starch accumulation, starch composition, stress response, and the phenylpropanoid pathway^56–58^. While the peak genetic markers identified by the FarmCPU resampling based GWAS were located in intergenic space between two of the tandem gene copies, a number of non-synonymous polymorphisms within coding sequences were in high linkage disequilibrium with the peak genetic markers. Four non-synonymous polymorphisms in perfect linkage -- consisting of a three base pair deletion, a three base pair insertion, and two point mutations -- within the first exon of Sobic.006G070564 where in tight linkage with the peak markers identified in the GWAS analysis, forming two distinct haplotypes at this locus. LefSE analysis of microbial taxa at the genus level, using haplotype at this position as the explanatory variable, identified significant changes (p-value<0.05) in the abundance of several genera from both subjects. Similar to the GWAS data, the abundance of *Escherichia* was higher in samples from both subjects fermented with the grain of sorghum varieties carrying the minor haplotype, and in both subjects the abundance of *Faecalibacterium* and other taxa (members of the *Ruminococcaceae* and *Lachnospiraceae*) was greater in samples fermented with the grain of sorghum varieties carrying major haplotype (Figure 5B). Total starch content was also significantly higher (p-value=0.0025, effect=2.04%) among sorghum varieties carrying the minor haplotype at this locus (Figure 5C), which is somewhat unexpected since the amylolytic bacterial taxa mapping here are more abundant in fermentations from lines carrying the major allele. However, CWIN are pleiotropic, affecting not only total starch, but also starch composition (amylopectin:amylose ratios) and constitutive expression of CWIN in maize can increase total starch while also reducing amylose content^59^. This matches well with our microbial phenotypes as *Faecalibacterium* and other amylolytic species of the *Ruminococcaceae* and *Lachnospiraceae* which preferentially ferment amylose and map to MEL6A are elevated in lines carrying the major allele. Consequently, one would hypothesize that the minor allele haplotype in Sobic006G070564 increases starch content in the seed but reduces its amylose content, having a major effect on the gut microbiome.

**Figure 5.**
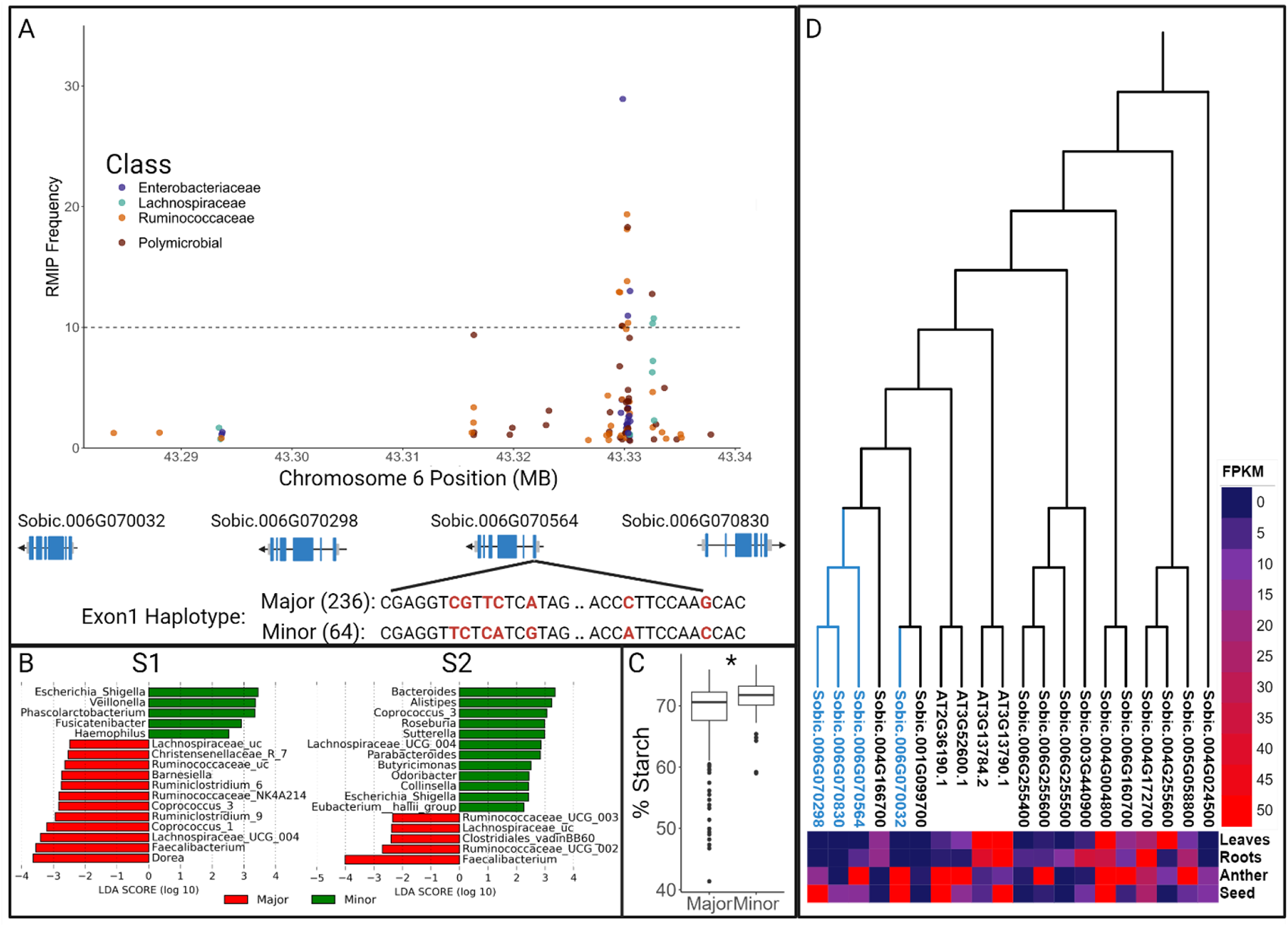
The SNPs most frequently associated microbiome metrics within MEL6A were within a tandem array of sorghum genes encoding cell wall invertases, four linked non-synonymous mutations are present in the first exon of Sobic.006G070564 (A). Presence of the mutations (Minor haplotype) is explanatory for the response of several microbes in both subjects (B) and seed starch content (C). These genes are highly expressed in the seed and are more similar in sequence to a well-studied tandem array of Arabidopsis cell wall invertases than other copies found in sorghum (D).

### Variation at the MEL6A locus produces consistent effects across diverse human microbiomes

The effect of tannins (MEL2 and MEL4B) and the haplotype group at MEL6A, drove strong and similar microbiome responses, including organisms that are members of the family *Lachnospireace* family and *Faecalibacterium* genus and we have previously characterized those responses^15^. Similar microbiome responses were also observed at MEL6A though there are no known genes that are part of the proanthocyanidin pathway within the MEL6A and indeed it seems that the phenotype could be driven by starch composition. To determine if the microbiome effects of tannin content (MEL2, MEL4B) and MEL6A are additive or due to a possible interaction relationship between loci, sorghum lines were pooled first by allele at MEL6A peak and then by presence/absence of tannins. Each pool was processed in the AiMs platform and screened across the microbiomes of twelve human donors where microbiomes from subject 1 and subject 2 were the same fecal collection as in the full panel mapping experiment (see Materials and Methods). Both the MEL6A haplotype allele and tannin content impacted microbial community structure in several subjects as indicated in an analysis of beta-diversity where significant differences by haplotype allele in ten of the twelve subjects were observed (SFigure 4). Across the top 30 genera present in all twelve human subjects based on 16S sequencing, several showed significant differences based on haplotype of pool (p-value<0.05). Although the effect of MEL6A haplotype was microbiome-dependent (e.g., different organisms were observed responding to the MEL6A haplotype effect in all subjects), the MEL6A haplotype nonetheless affected multiple taxa in each microbiome (Figure 6A, B). In an analysis of the average log2 fold change in all subjects the response of many taxa was as expected from the initial analysis, with higher abundances of *Anerostipes*, *Faecalibacterium*, *Coprococcus*, and *Dorea* in lines and pools that contain the major haplotype, and lower abundances of *Escherichia* (Figure 6C). The response of many genera was consistent across experiments even when compiling subjects in the log2 fold change analysis, (Pearson correlation, R=.39, p-value=0.02). In three of twelve subjects’ microbiomes the abundance of *Faecalibacterium* was significantly higher in the MEL6A major allele haplotype pools (Figure 6A and 6B). To further investigate the effect of the MEL6A haplotype on *Faecalibacterium*, the absolute abundance of *Faecalibacterium* was quantified using qPCR for microbiomes from each of the twelve subjects treated with the haplotype pools. With this more quantitative measure, the amount of *Faecalibacterium* was significantly higher in microbiomes treated with the minor allele in eight of the twelve subjects (Figure 6D and E). In an analysis of variance (ANOVA) of absolute abundance of *Faecalibacterium* quantified by qPCR, with factors subject, presence/absence of tannins, allele at MEL6A, and all interactions between them, all factors were highly significant (p-value<0.0005) except the interaction term for tannin and MEL6A allele (Table S8). Indicating while the allele at MEL6A and the effects of allelic variation at MEL2 and MEL4B on tannin content both affect growth of *Faecalibacterium*, they appear to do so independently, and thus likely affect this genus through variation in different components of the seed (e.g., tannins and starch composition).

**Figure 6.**
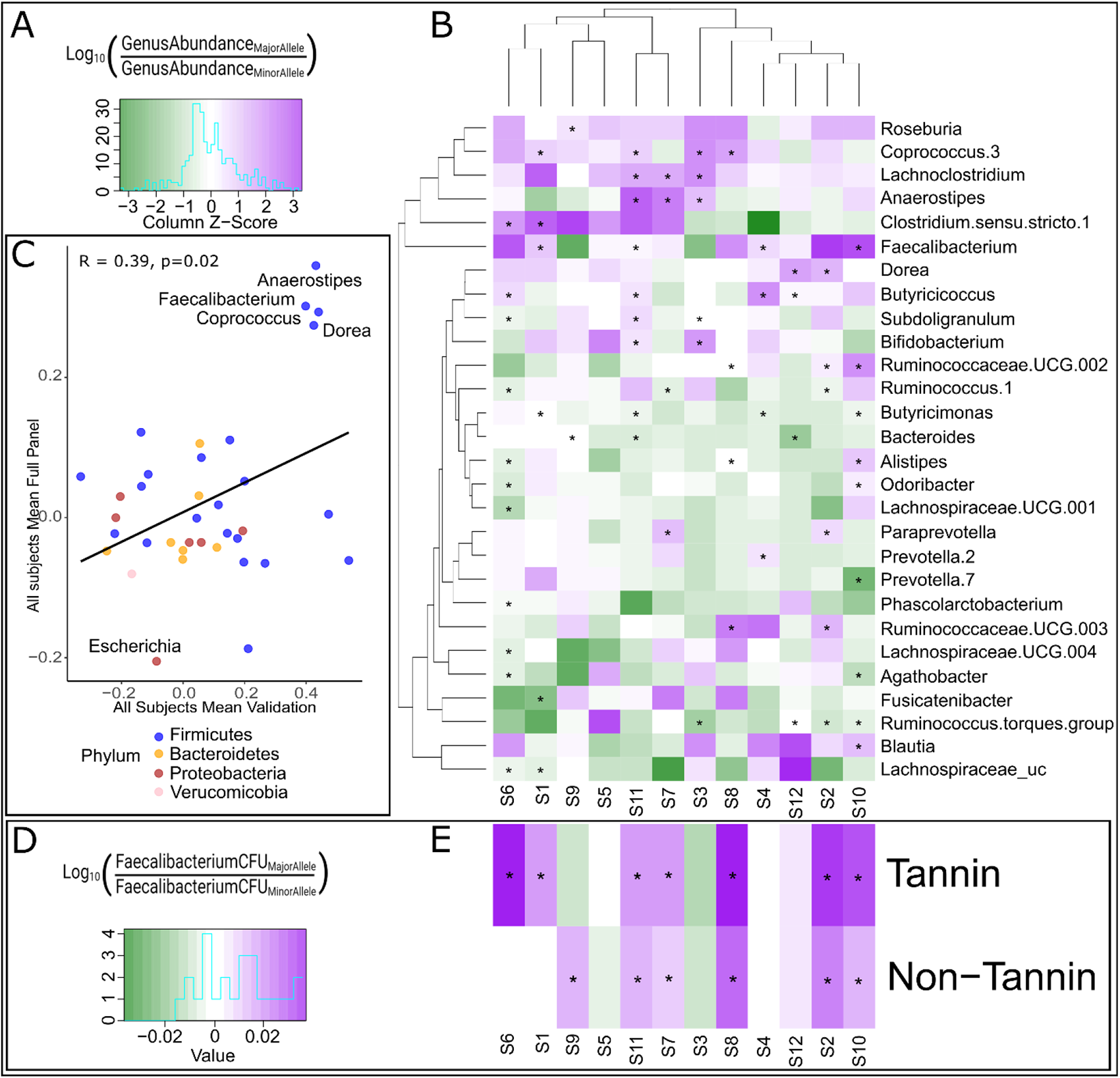
The equation and heatmap legend (A) for log ratios of top genera from the microbiomes of 12 human subjects treated with the major haplotype by minor haplotype at the MEL6A allele within the tannin containing pools (B). Scatter plot of log2 fold change in genera in AiM reactions of major haplotype/minor haplotype lines (x-axis) and pools (y-axis), each genus value is the average fold change of all subjects in the respective experiment (n=2 for full panel, n=12 for validation) and is colored according to its phyla. R and p-value based on Pearson correlation (C). The equation and heatmap legend (D) for the mean values of Log ratios of *Faecalibacterium* from the microbiomes of 12 human subjects quantified by qPCR treated with the major haplotype by minor haplotype at the MEL6A allele in both tannin and non-tannin pools (E). Asterisks denote significant p-value from the result of a Wilcoxon test p<0.05.

## Discussion

### Repeatability of mapping data across multiple populations

In this study, we identified a total of 15 MELs distributed across the sorghum genome where sorghum genetic variation produces substantial impacts on diverse human gut microbiomes. This included reidentification and more precise localization of four out of five broad QTL loci described in a previous study conducted using a biparental sorghum mapping population^15^. The increased number of MELs identified is consistent with the better presentation of genetic and phenotypic diversity provided by association panels relative to recombinant inbred populations, and the more precise localization is consistent with the larger number of recombination events captured by association mapping relative to structured populations such as recombinant inbred populations. At the same time, the reidentification of 80% of the MELs described in a study conducted using a different sorghum population grown in a different environment – Yang et al.^15^ used grain from individual plants grown in greenhouses, our study uses grain from small plot trials grown in the field -- suggests that the MELs we identify are likely to generalize across sorghum populations and provide consistent effects on human gut microbiomes when sorghum is grown in different parts of the globe and in different years, increasing the feasibility of breeding sorghum for beneficial microbial outcomes. Of the 15 MEL from the current study, four (MEL2, MEL3A, MEL4B, MEL6B) correspond to four of the five total MEL detected in the Yang et al.^15^ study, which used RIL population having much less genetic diversity than the SAP. Thus, genetic diversity within the SAP not only validated previous findings, it also further extended the catalogue of loci in *Sorghum bicolor* where genetic variation has multiple effects on the microbiome. In both studies, tannin content and microbes belonging to the families *Ruminococcaceae* and *Lachnospiraceae* mapped with overlapping peaks to MEL2 and MEL4B, which correspond to the Tan 2 and Tan 1 regulatory genes, respectively, which control synthesis of condensed tannins. Remarkably, though we used microbiomes from different subjects in the two studies, the microbiome phenotypes of MEL2 and MEL4B across multiple human subjects consistently showed major effects of variation on *Faecalibacterium* and other members of the *Ruminococcaceae* along with members of the *Lachnospiraceae*. Thus, even in very distinct genetic backgrounds (SAP vs BTx623 X IS3620C RILs), variation at Tan1 and Tan2 imparts conserved signature on the human gut microbiome across diverse human subjects.

### The human gut microbiome as a diverse set of phenotypes to discover novel seed traits

In addition to validating previous findings, it is worth noting that our emphasis on MEL, which affect multiple microbial taxa from different microbiomes, focuses the genetic analysis on loci where variation can have a major effect across human populations. This was apparent from the similar effect of MEL6A on the absolute abundance of *Faecalibacterium* (Figure 6D and E) and members of the *Lachnospiraceae* across 10 different human subjects (Figure 6A). Despite the individuality of the gut microbiome in humans, emphasis on MEL enables us to use relatively small numbers of microbiomes for mapping across large numbers of sorghum genotypes, ultimately allowing us to survey a significant proportion of genetic variation present within the crop. The 11 new MEL identified in this study also had significant associations with multiple organisms and/or polymicrobial traits from the microbiomes of both subjects used in the phenotyping and we fully expect genetic variation at these loci to also have major effects across microbiomes of additional human subjects. We observed many overlapping peaks for microbial groups at different levels of taxonomic resolution and polymicrobial traits, however in some instances, polymicrobial traits mapped at loci where no single taxa mapped uncovering loci associated with relationships within the microbiome (Figure S5).

### Co-localization of microbiome traits and agronomic/biochemical traits

In addition to multiple microbiome traits mapping to each MEL, we also found that multiple agronomic, seed, or biochemical traits also mapped within each of the MEL. Interestingly, there was only slight enrichment for biochemical or seed traits as roughly 21.7% and 22.5% respectively of significantly associated markers are within MEL compared to 15.9% markers associated with agronomic traits. However, as we illustrated at MEL6A, the co-localized microbiome and seed traits, along with known genetic variation at the locus, facilitated hypothesis generation about causal variants and mechanism through which such variation impacts the microbiome. We note that significant associations for traits associated with seed color (tannins, polyphenols, etc), seed size/weight, and the content of seed starch, total fiber, and oil content co-localized along with microbiome phenotypes to nearly every MEL, apart from MEL3A (Table S7). As is illustrated with the known pleiotropic effects of variation at Tan 1 on tannins, seed color, starch content, and microbiome^15^, genetic variation driving phenotypes corresponding to the MEL can be quite pleiotropic.

### Candidate genes and hypothesis generation for effects of MEL on the microbiome

Co-localization of seed traits and microbiome traits at a MEL can help drive hypotheses for candidate genes/pathways that drive the seed and microbial phenotypes. Beyond the Tan 1 and Tan 2 loci, we also note the tandem array of cell wall invertases (CWIN) found at MEL6A, whose β-fructofuranosidase activity (sucrose → glucose + fructose) can influence a wide array of plant functions such as growth and development, carbon/nutrient partitioning, and responses to biotic and abiotic stress. Thus, if the CWIN haplotype associated with the MAF in MEL6A is responsible for the microbiome and seed traits, it may be manifested through a number of mechanisms. We initially tested total fructans across seed with the major vs minor allele and CWIN haplotype content at MEL6A and found no statistically significant differences (data not shown). We did, however, detect significant effects on total starch content and many of the microbial taxa mapping to MEL6A are known amylolytic organisms who would be expected to show differences if starch content were driving the microbial phenotype. More work is necessary to determine if it is composition of the starch (e.g. amylose:amylopectin ratios) is driving the phenotype, but it is clear that expression of CWIN in maize can drive very similar starch phenotypes (elevated total starch but with decreased amylose content) that would be expected to drive the microbial phenotypes that are associated with MEL6A.

### The human gut microbiome as an agnostic approach for identification of traits in food crops that can affect human health and nutrition

While we can currently only generate mechanistic hypotheses around a small number of the MEL, the nature of the MEL themselves (major effects on multiple microbes and the ability to detect allelic effects of MEL across different human subjects) illustrates the strength of this approach for identifying novel plant traits affecting human health and nutrition. Unlike mapping specific biochemical traits, use of different features of the human gut microbiome as a complex combination of traits (e.g., abundances of individual taxa, PCs comprising multiple taxa, prebiotic indices) allows us to agnostically interrogate the effects of plant genetic variation on a wide variety of traits that can have a major impact on human health. Moreover, once MEL are discovered, one can simply reduce the phenotypic analysis to individual microbes, groups of microbes, microbial metabolites (e.g., SCFA) to simplify phenotypic analysis. While the exact biochemical pathway that drives a MEL may not be known, the microbial trait itself (through AiMS-based fermentations) can be used in an efficient and high-throughput manner to facilitate further genetic analysis to localize causal variants and downstream breeding to improve the trait.

## Supporting information

Supplemental Dataset 1, BLUES calculated for all microbiome metrics and sorghum agrinomic traits.

Supplemental Dataset 2, Compiled GWAS output from all traits.

Supplemental Dataset 3, Faecalibacterium qPCR data from Validation study.

Supplemental Table 1, Microbial genera that comprise the "Prebiotic Index".

Supplemental Table 2, List of sorghum genotypes used in study.

Supplemental Table 3. Description of agrinomic traits used in study.

Supplemental Table 4. Heritability values for microbiome genera calculated in small scale experiment with microbiomes from eight human subjects.

Supplemental Table 5, Heritability values for microbiome genera calculated in large-scale mapping experiment.

Supplemental Table 6, Description of LD calculations for candidate loci.

Supplemental Table 7, Description of all MELs identified by RMIP GWAS.

Supplemental Table 8. Results of ANNOVA analysis from Faeclaibacterium qPCR data from validation study.

Supplemental Figure 1, Correlation of prebiotic index to butyrate.

Supplemental Figure 2, Genus level abundance of microbiomes from two subjects used in mapping experiment.

Supplemental Figure 3, LD analysis of associated SNPs in each MEL.

Supplemental Figure 4, Beta-diversity (Weighted Jaccard index) analysis of 12 human microbiomes screened with sorghum lines pooled by allele at MEL6A.

Supplemental Figure 5, GWAS output for Faecalibacterium at ASV and genus level, the latent variable and principal component with the most associated m

## Data Availability

The DNA sequencing reads for this study are available in the NCBI SRA database as project accession PRJNA1012736. All ASVs were assigned with taxonomic information using the taxonomy classifier SILVA database^28^. Best linear unbiased estimates (BLUEs) calculated for microbiome metrics and sorghum agronomic data are included in Supplemental Dataset 1. Complete GWAS output of all traits is included in Supplemental Dataset 2. The qPCR output of *Faecalibacterium* in the MEL6A validation study are included in Supplemental Dataset 3.

