## Supplementary figures and images for "Grain Utilization by the Gut Microbiome as a Human Health Phenotype to Identify Multiple Effect Loci in Genome-Wide Association Studies of *Sorghum bicolor*"

### Supplemental Figure 1, Correlation of prebiotic index to butyrate.

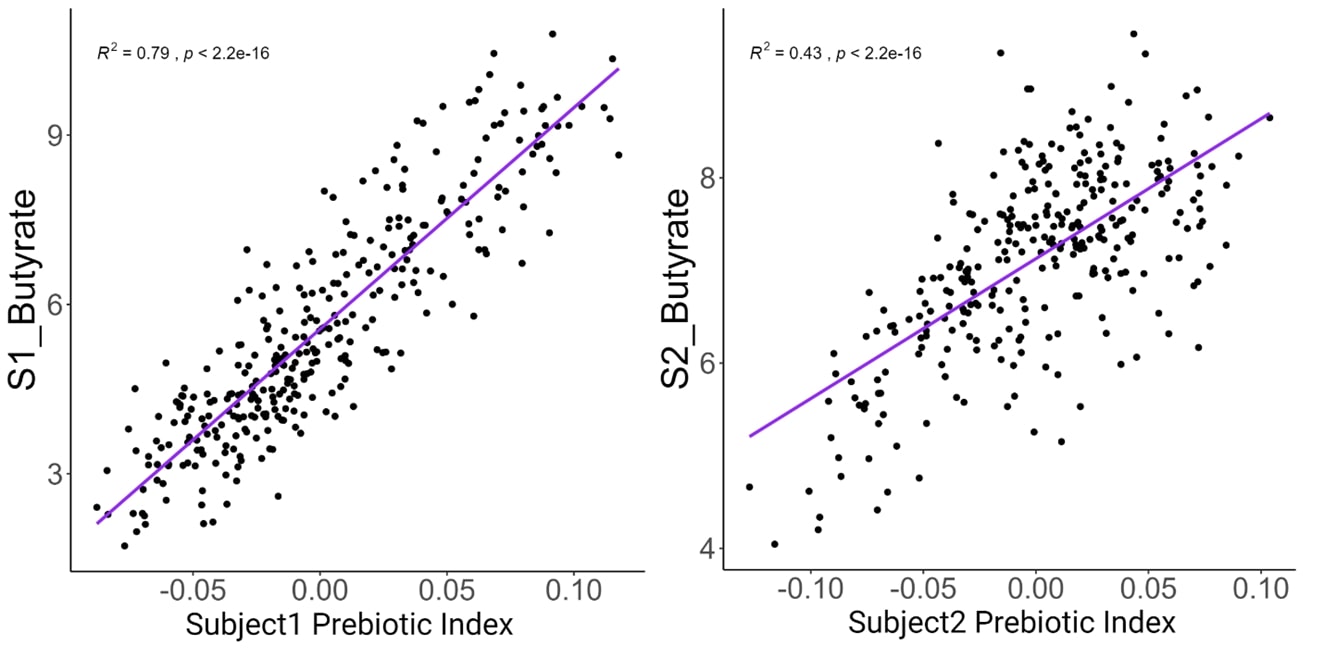

### Supplemental Figure 2, Genus level abundance of microbiomes from two subjects used in mapping experiment.

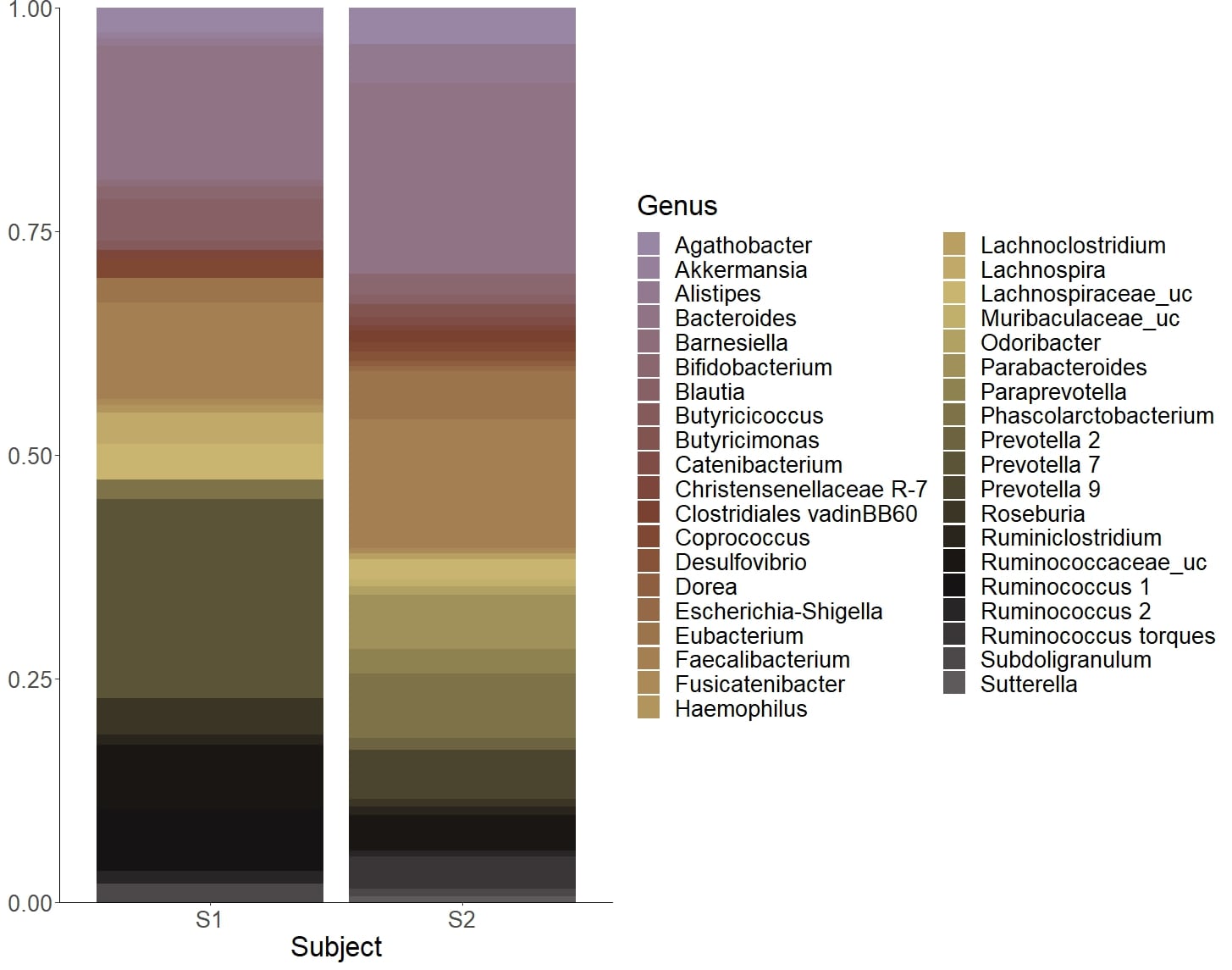

### Supplemental Figure 3, LD analysis of associated SNPs in each MEL.

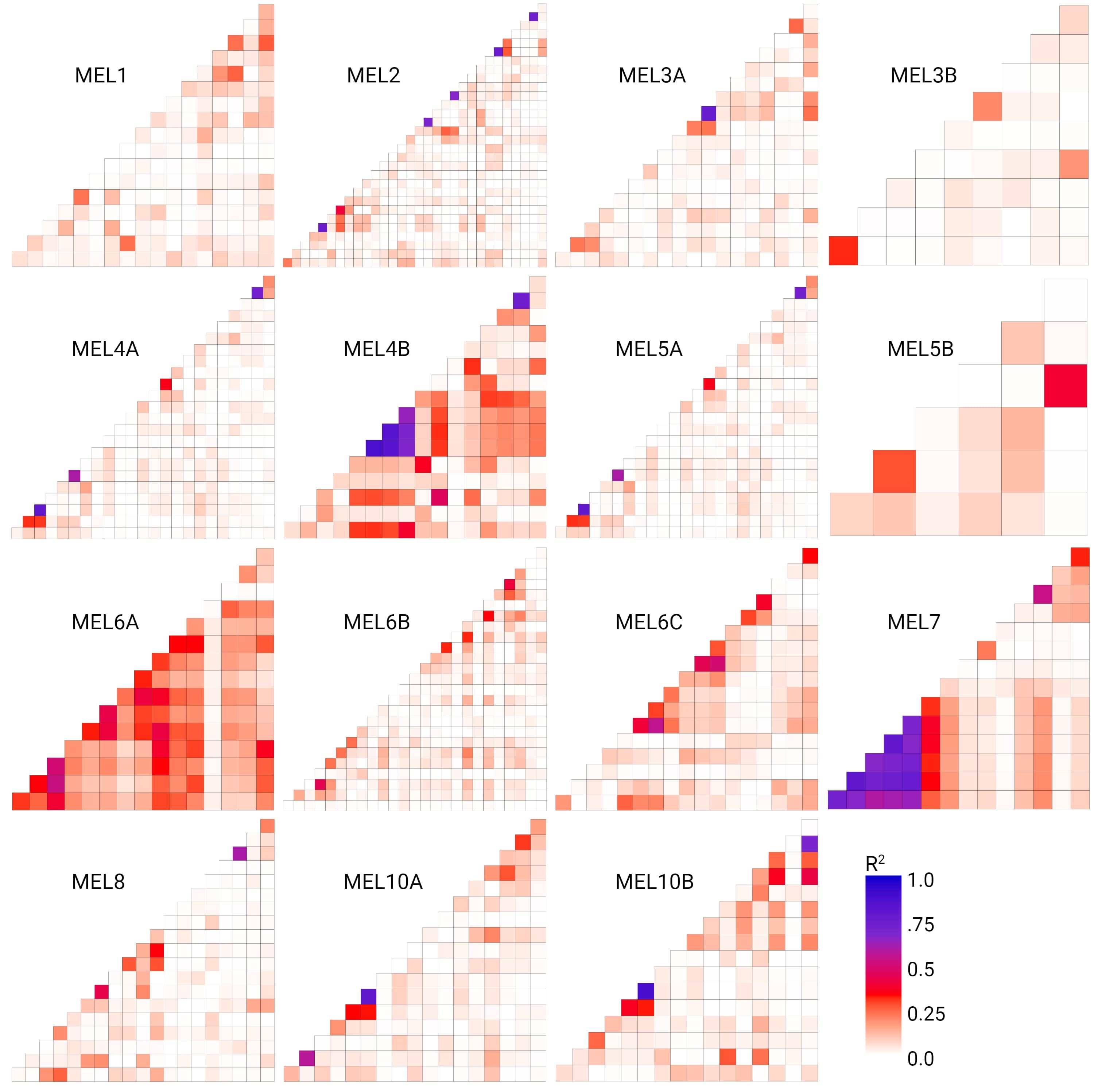

### Supplemental Figure 4, Beta-diversity (Weighted Jaccard index) analysis of 12 human microbiomes screened with sorghum lines pooled by allele at MEL6A.

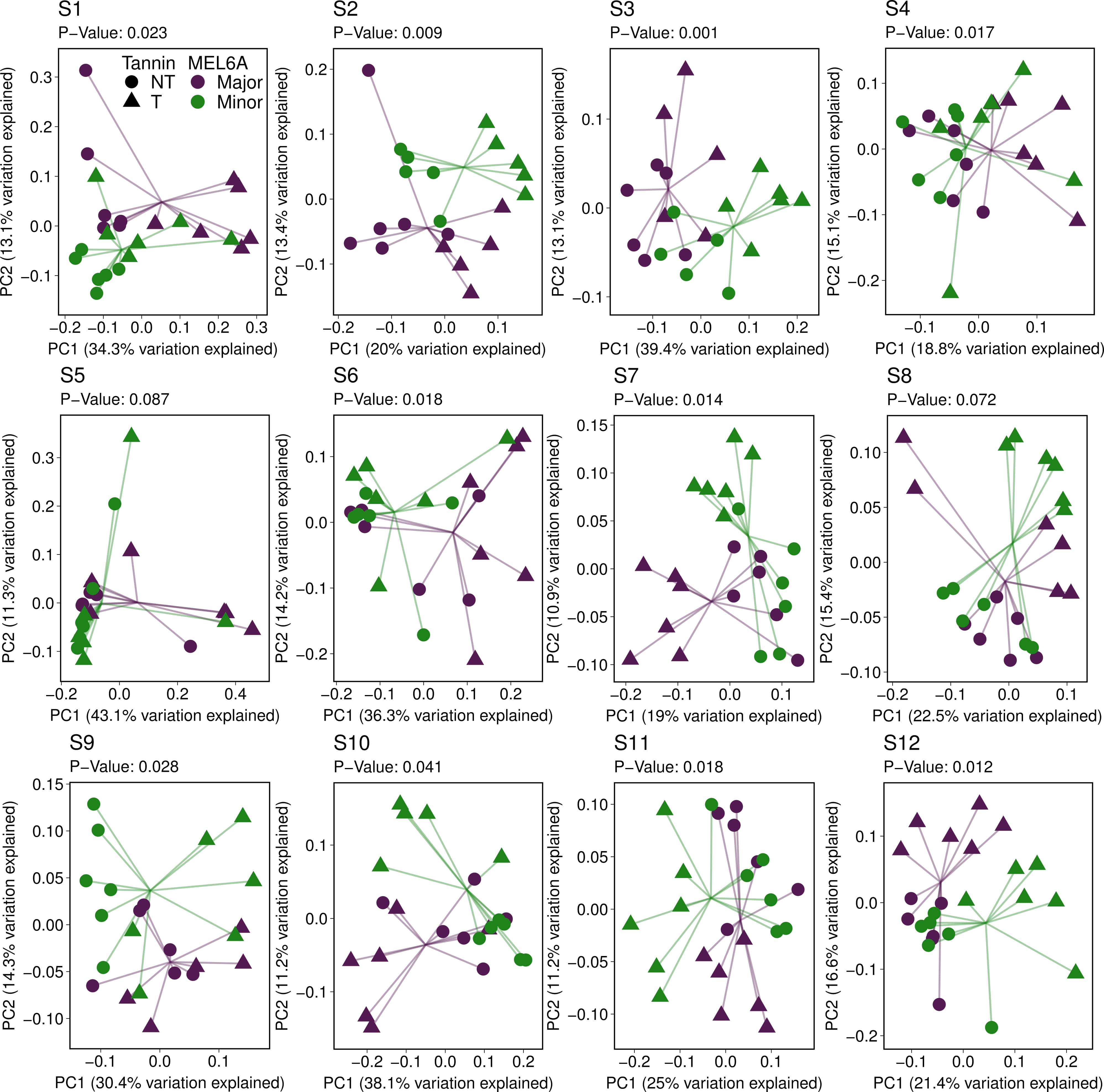

### Supplemental Figure 5, GWAS output for Faecalibacterium at ASV and genus level, the latent variable and principal component with the most associated m

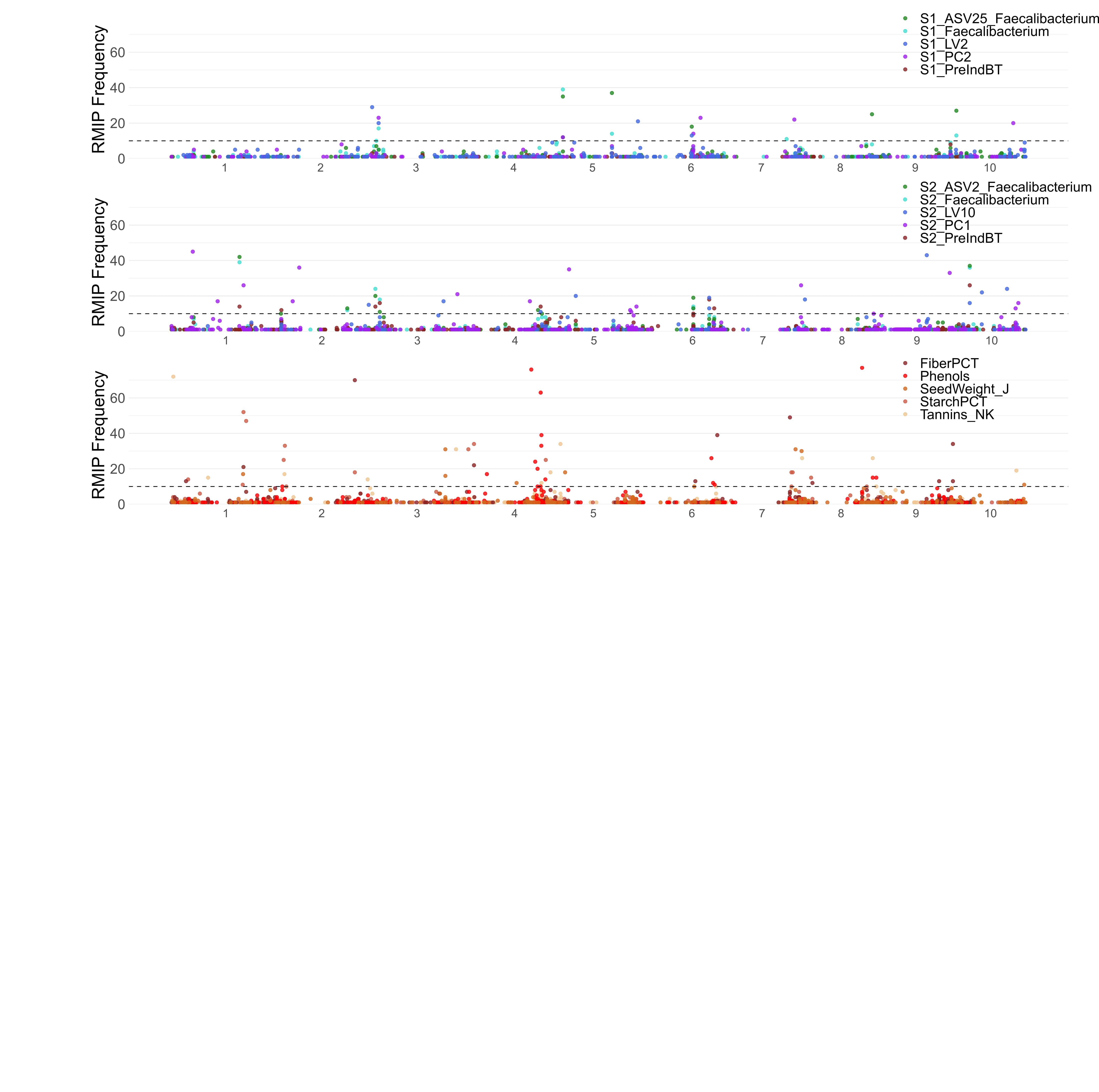
